# Improving peptide-level mass spectrometry analysis via double competition

**DOI:** 10.1101/2022.05.11.491571

**Authors:** Andy Lin, Temana Short, William Stafford Noble, Uri Keich

## Abstract

The analysis of shotgun proteomics data often involves generating lists of inferred peptide-spectrum matches (PSMs) and/or of peptides. The canonical approach for generating these discovery lists is by controlling the false discovery rate (FDR), most commonly through target-decoy competition (TDC). At the PSM level, TDC is implemented by competing each spectrum’s best-scoring target (real) peptide match with its best match against a decoy database. This PSM-level procedure can be adapted to the peptide level by selecting the top-scoring PSM per peptide prior to FDR estimation. Here we first highlight and empirically augment a little-known previous work by He et al., which showed that TDC-based PSM-level FDR estimates can be liberally biased. We thus propose that researchers instead focus on peptide-level analysis. We then investigate three ways to carry out peptide-level TDC and show that the most common method (“PSM-only”) offers the lowest statistical power in practice. An alternative approach that carries out a double competition, first at the PSM and then at the peptide level (“PSM-and-peptide”), is the most powerful method, yielding an average increase of 17% more discovered peptides at a 1% FDR threshold relative to the PSM-only method.

## 1 Introduction

The goal of most proteomics tandem mass spectrometry experiments is to detect and quantify the proteins present in a complex biological sample. Discoveries from such an experiment are most useful when they are associated with a statistical confidence estimate. Such estimates are typically provided by thresholding with respect to a specified *false discovery rate* (FDR), defined as the expected rate of false discoveries among the set of all accepted discoveries.

In practice, this type of FDR control can be carried out at one of three different levels, corresponding to three types of discoveries. First is the peptide-spectrum match (PSM) level, where each discovery is an observed spectrum linked to the peptide that is inferred to be responsible for generating the spectrum. Second is the peptide level, in which multiple PSMs for the same peptide sequence are considered jointly. Third is the protein level, where evidence is accrued across all peptides associated with a given protein. The decision of whether to control the FDR at the level of PSMs, peptides, or proteins depends upon the question at hand, and it is not unusual for a single study to employ more than one type of FDR control for different purposes.

In mass spectrometry proteomics, the most widely used methods for FDR control are based on a straightforward procedure known as *target-decoy competition* (TDC).^1^ The procedure estimates and consequently controls FDR at the PSM level by comparing scores from a search of each observed spectrum against a database of real (*target*) peptides with scores for the same spectrum against a database of reversed or shuffled *decoy* peptides. This method is called target-decoy *competition* because each spectrum only receives a single score: the target and decoy peptide matches to the spectrum compete against one another, and the highest scoring match is assigned to it.

The TDC procedure relies on the underlying assumption that an incorrect match (i.e., a false discovery) is equally like to have come from a target or a decoy peptide. Hence, the number of false discoveries above some score threshold can be estimated by the number of decoys that win the competition and whose scores are above the same threshold. Thus, dividing this estimated number of false discoveries by the number of target peptide wins that score above the same threshold yields an estimate of the FDR. He et al. showed that if we further assume that the scores of the incorrect matches are independent of one another (as well as of the correct matches), and provided we add 1 to the number of decoy wins above the threshold before dividing by the number of target wins, then we can control the FDR at level *α* by choosing the smallest score threshold for which the estimated FDR is still ≤ *α*.^2^

Partly due to its simplicity to understand and implement, TDC is by far the most widely used method for PSM-level FDR control in proteomics mass spectrometry. Recently, the same competition-based approach to control the FDR gained significant popularity in the statistics and machine learning communities after Barber and Candès introduced their knockoff filter.^3^

Motivated by the PSM-level TDC, decoy-based approaches to controlling the FDR at the peptide and protein levels have also been described. In both cases the idea is to aggregate the scores of all PSMs involving a peptide (taking the maximum of the scores) or a protein (taking the sum of the peptide scores). We thus assign a score to each target and decoy peptide (or protein, depending on the level of our analysis). With these scores in hand we can continue analogously to the PSM-level TDC, although as we will see below there is more than one way to implement how the scores are aggregated and how the competition is implemented.

Our goal in this paper is to convey two distinct messages related to decoy-based FDR control in mass spectrometry proteomics. The first is essentially reiterating the message of He et al. that controlling the FDR at the level of PSMs is problematic, due to unavoidable potential dependencies between the incorrect PSMs. As we mentioned above, TDC provably controls the FDR, but only under certain assumptions. However, as He et al. have pointed out, in practice these assumptions might not be met.^2^ Here we provide further empirical evidence showing that PSM-level FDR estimates can be liberally biased, meaning that if you try to control the FDR at, say, 1%, then the TDC procedure is likely to control the FDR at a less stringent threshold. In our experiments, the magnitude of the effect becomes larger as the FDR threshold increases.

Our second message is that the most common way to control FDR at the peptide level is suboptimal, in the sense that it yields fewer detected peptides than a relatively simple variant of that procedure. Though it is rare in the literature to precisely specify how peptide-level FDR is performed, the few cases where we found a precise description appear to carry out PSM-level competition only.^2,4,5^ Specifically, the explicit pairing between each target peptide and its shuffled decoy is ignored. Instead, each peptide (target or decoy) is assigned a score which is the maximum scoring PSM involving that peptide. Note that this is equivalent to considering the weeded-out list of PSMs obtained by removing PSMs that are matched to a peptide that has a higher scoring PSM associated with it. The FDR is next estimated and controlled using the number of decoy peptides (or weeded-out decoy PSMs) above the threshold, as noted above for the PSM-level TDC.

Here we investigate two additional variant techniques for carrying out peptide-level FDR control using TDC and compare all three procedures. These variants were motivated in part by the “picked protein” approach for protein-level FDR control.^6^ The idea, whose benefits were demonstrated in^7^ and,^8^ is to perform a “head-to-head” competition between each target protein and its paired decoy, keeping only the higher scoring of the two. Thus, we compared the above PSM-only competition approach to peptide-level analysis with two additional protocols, both of which involve a direct peptide-level competition between each target peptide and its paired decoy (so only the higher scoring of the two is kept). The first protocol, peptide-only, involves only the peptide-level competition, i.e., the PSMs are generated by separately searching the target and decoy databases. The second protocol, PSM-and-peptide, involves a double competition. First, the PSMs are generated by searching a concatenated database. This first competition is the same as in the commonly used approach, but in PSM-and-peptide it is followed by a second, peptide-level, competition. After generating the PSMs and performing the peptide-level competition both peptide-only and PSM-and-peptide proceed as in the PSM-level TDC.

We find that PSM-and-peptide, which uses competition first at the PSM level and then at the peptide level, yields the greatest number of discovered peptides, whereas the commonly used PSM-only competition yields significantly lower statistical power. Notably, switching from PSM-only to PSM-and-peptide is relatively straightforward, as long as it is possible to match each target peptide to its corresponding decoy.

## 2 Methods

### 2.1 Methods for FDR control

#### 2.1.1 PSM-level TDC

The goal of TDC carried out at the PSM level is to estimate and control the false discovery rate among a collection of PSMs produced by a database search engine. We are given a set *S* of *n* spectra, and we assume that the spectra have already been searched against a target database 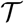 and a decoy database 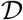. For each spectrum we retain the top-scoring PSM, breaking any ties randomly. We refer to the scores of the PSMs that involve target peptides as *t*_1_,*t*_2_, …, *t_m_t__* and the scores of decoy PSMs as *d*_1_, *d*_2_, …, *d_m_d__*, where *m_t_* + *m_d_* = *n*. We can then estimate the FDR among all PSMs that score greater than a specified score threshold *τ* (assuming that larger scores are better) as

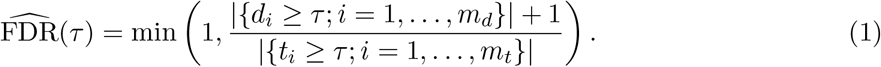

Intuitively, the denominator represents the number of discoveries of interest (the target PSMs), and the numerator is our decoy-based estimate of the number of false positives among those discoveries.

In practice, rather than specifying a priori a score threshold *τ*, we typically specify the desired FDR threshold *α* and choose our rejection threshold, *τ*(*α*), as the smallest *τ* for which the estimated FDR is ≤ *α*:

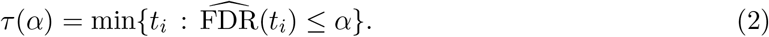

The estimated FDR in Equation (1) differs from the one offered by Elias and Gygi^1^ in three ways. First, their formulation includes a factor of 2 in the numerator and includes both target and decoy PSMs in the denominator. This approach thus controls the FDR among the combined set of target and decoy PSMs. In practice, it typically is of more interest to control the FDR only among the targets, as in Equation (1). Second, the numerator in our formulation includes a +1 that is missing from the Elias and Gygi formulation. This +1 correction is required in order to achieve valid FDR control.^2,9^ In practice, this +1 correction will have a negligible effect except in the presence of very few discoveries or a very stringent FDR threshold. Finally, our formulation includes an enclosing min operation, which simply ensures that we do not report an estimated FDR > 1.

#### 2.1.2 PSM-only competition for peptide-level FDR control

The most commonly used method for estimating FDR at the peptide level, which we refer to as “PSM-only” (Figure 1B, Supplementary Algorithm 1) is quite straightforward.^4^ This procedure starts by carrying out PSM-level target-decoy competition, retaining only the top-scoring PSM per peptide sequence. Thereafter, the FDR among the target peptides is estimated and controlled using the analogs of Equations (1) and (2), where the scores *t_i_* and *d_i_* refer to the peptide rather than PSM scores.

**Figure 1:**
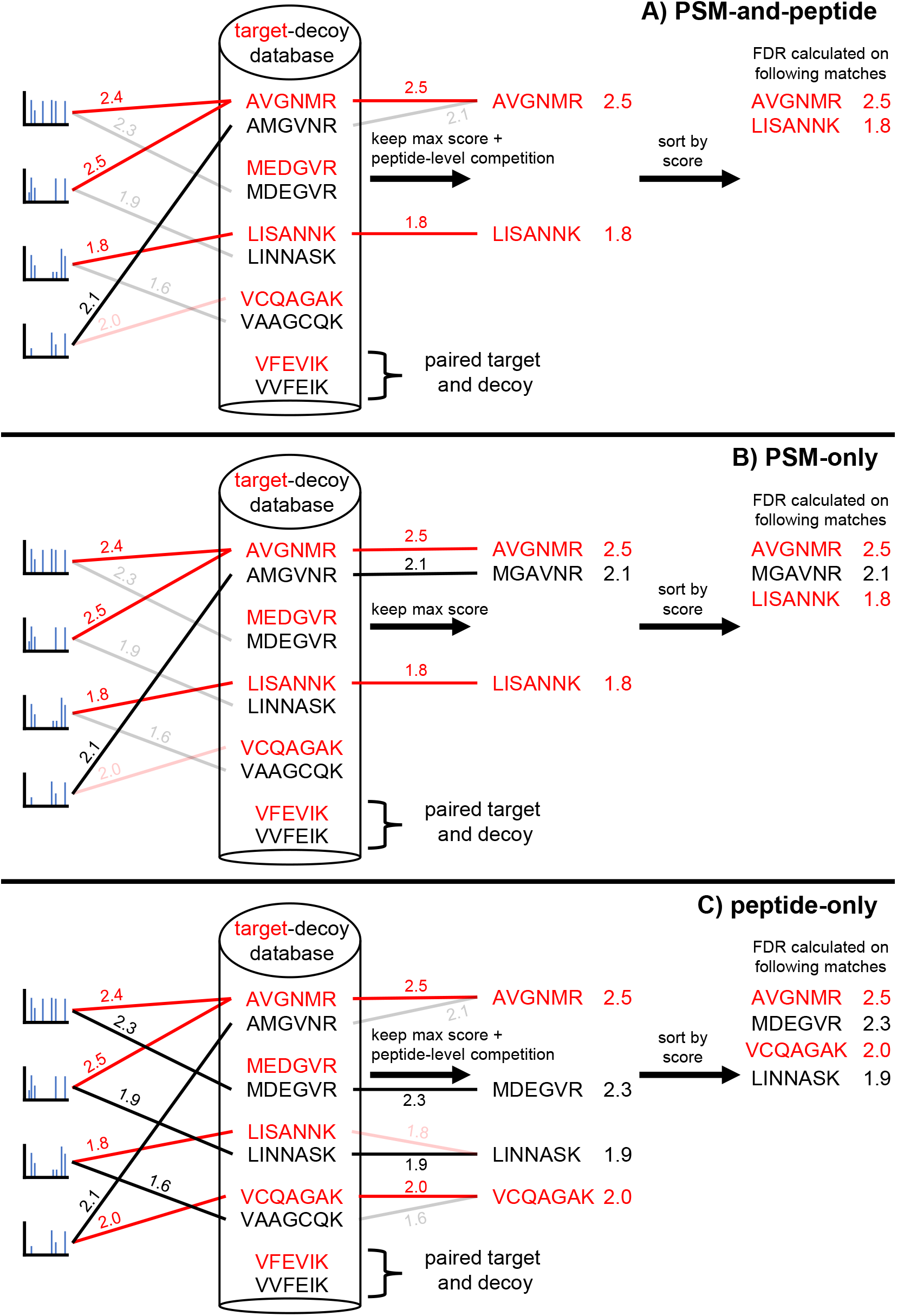
Graphical view of peptide-level estimation methods. Graphical view of three different procedures for estimating peptide-level FDR: A) PSM-and-peptide, B) PSM-only, and C) peptide-only.

#### 2.1.3 Peptide-only and PSM-and-peptide competition for peptide-level FDR control

The next two procedures for peptide-level FDR control procedure mimic a crucial step of the “picked protein” procedure for protein-level FDR control.^6^ Specifically, both the “peptide-only” (Figure 1C, Supplementary Algorithm 2) and “PSM-and-peptide” (Figure 1A, Supplementary Algorithm 3) procedures employ a head-to-head competition between each target peptide and its paired decoy so that only the higher scoring of the two is kept.

The difference between the two methods is that the PSM-and-peptide approach creates PSMs by scanning each spectrum against the concatenated target-decoy database, keeping only the best matching peptide for the scanned spectrum. In contrast, the peptide-only approach does not employ a PSM-level competition; instead, it separately scans each spectrum against the target and the decoy databases, keeping the corresponding two best matches for the scanned spectrum.

Both methods assign each target or decoy peptide a score that is the maximum of all PSMs involving the peptide. Both methods then carry out peptide-level competition for each target-decoy peptide pair. The rest follows exactly as in the PSM-only approach, where the FDR among the target peptides is estimated and controlled using Equations (1) and (2), where the scores refer to the peptide scores.

### 2.2 Datasets

To evaluate our various FDR control methods, we used runs from six different mass spectrometry datasets. These six datasets consisted of data from the ISB18 mix,^10^ castor plant,^11^ *E. coli*,^12^ human,^12^ mouse,^13^ and yeast samples.^14^ All nine runs from the ISB18 dataset were downloaded from http://regis-web.systemsbiology.net/PublicDatasets. Two runs from each of the five other datasets were downloaded from PRIDE^15^ (Table 1). Raw mass spectrometry runs were converted to MS2 format using MSConvert version 3.0.^16^ The protein sequence databases corresponding to the castor plant, *E. coli*, human, mouse, and yeast proteomes were downloaded from Uniprot (www.uniprot.org) in October 2021 or January 2022. The sequences corresponding to the ISB18 mix were downloaded from https://regis-web.systemsbiology.net/PublicDatasets/database.

**Table 1:**
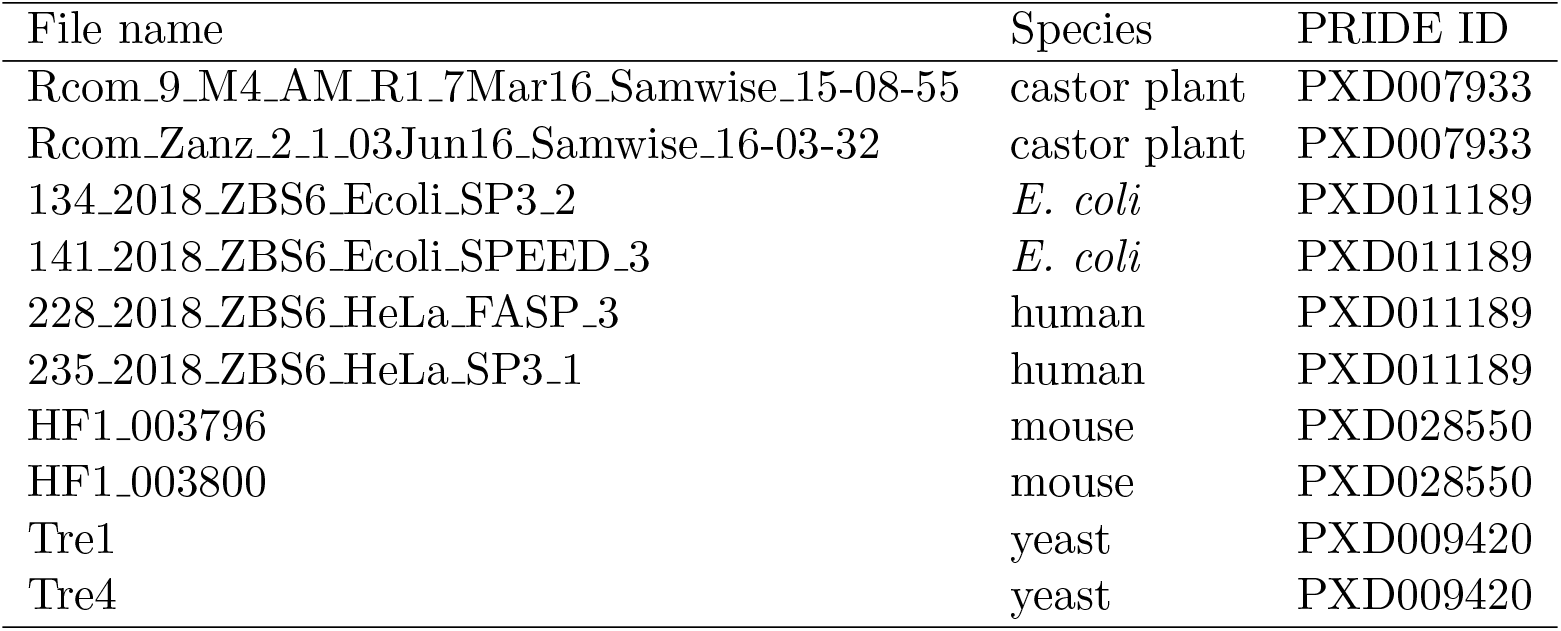
Datasets used in this study.

### 2.3 Database searching

When comparing the power of the procedures we used the Tide search engine,^17^ as implemented in Crux version 4.1,^18,19^ to search the aforementioned datasets of spectra from the castor plant, *E. coli*, human, mouse, and yeast runs against a target-decoy database containing the proteome of the sample being analyzed. The respective target sequences were obtained as described above while decoy sequences were generated by the tide-index tool in Crux by shuffling each target peptide sequence, leaving the N-terminal and C-terminal amino acids in place.

Each file was searched using four different score functions: XCorr,^20^ XCorr p-value,^21^ combined p-value,^22^ and Tailor.^23^ The precursor mass tolerances were estimated using Param-Medic^24^ and set to be 85, 40, 35, 40, and 25 ppm for the castor plant, *E. coli*, human, mouse, and yeast runs, respectively. For these searches, all other parameters were set to their default values, except that `--top-match=1` and one missed cleavage was allowed. For the XCorr p-value and combined p-value score functions, `--exact-p-value=T` and `--mz-bin-width=1.0005079`. For the Tailor score function `--use-tailor-calibration=T`. We note that `--concat=F` by default, and therefore target and decoy results are reported separately.

### 2.4 Entrapment experiment

To evaluate whether each FDR estimation method properly controls the FDR we performed an entrapment experiment using the ISB18 dataset. In such an experiment, spectra are searched against a database containing the sequences in the sample in addition to a set of sequences not present in the sample.^25^ That is, the target database is concatenated with a set of additional, so-called entrapment sequences that are presumably not present in the sample being analyzed. Typically those sequences are from another proteome, and they should not be confused with the decoy sequences: if TDC is used then decoy sequences are constructed for the entire target database, including the entrapment sequences.

Importantly, the number of entrapment sequences is set to be much larger than the number of sequences that are present in the sample. In this case, the target database consisted of the ISB18 proteins augmented by the castor plant proteome providing the entrapment sequences. This yielded over 1,250 entrapment peptides for every potential in-sample ISB18 peptide. Since an incorrect identification is equally likely to involve any peptide in the combined database, the large ratio of entrapment-to-relevant peptides suggests that the vast majority of the false discoveries will involve entrapment peptides, and hence we can account for those. By the same token any match to the in-sample part of the database is very likely to be correct. This procedure allows us to reliably estimate the false discovery proportion (FDP) among the set of reported target PSMs and compare that FDP to the selected FDR threshold. Because the FDR is defined as the expectation of the FDP, this empirical FDP should not, on average, significantly exceed the FDR if the FDR estimation method is valid.

In this study, all scans from the nine ISB18 runs were collectively searched against a database containing the ISB18 proteins and the castor plant proteome. After removing peptide sequences in common, the castor plant and ISB18 proteomes contained 571,319 and 449 peptides, respectively. The FDP in the reported list of target PSMs was estimated by dividing the number of presumed incorrect PSMs (the ISB18 spectra that matched a castor plant entrapment peptide) by the total number of reported target discoveries (the ISB18 spectra that were matched with any part of the target database and scored above the cutoff).

The ISB18 spectra were searched using Tide against 10 different randomly shuffled decoy databases, where each decoy database was generated in Crux by shuffling the entrapment-augmented target database with a different random integer seed ranging from zero to nine, inclusive. All searches were done using the exact p-value score function of Tide with the precursor mass tolerances set to 50 ppm, `--exact-p-value=T`, and `--mz-bin-width=1.0005079`.

## 3 Results

### 3.1 PSM-level FDR control with TDC can be liberally biased

As mentioned above, in order to guarantee that TDC controls the PSM-level FDR, we need to assume that the incorrect PSMs are independent of each other. However, in practice we expect spectra that are generated from the same peptide to be highly correlated. Dynamic exclusion helps to reduce the magnitude of this problem but does not remove the problem, especially if, as is often done, *multiple runs are combined* before applying TDC.

The problem with such highly correlated spectra is that their corresponding optimal PSMs tend to involve the same peptide. If this match happens to be an incorrect match to a target peptide, then it can significantly inflate the actual FDP in a way that TDC cannot account for. In such cases, TDC may fail to control the FDR. Indeed, borrowing from He et al.,^2^ imagine an extreme case where the dataset consists of 100 spectra that were all generated by the same peptide and this peptide does not appear in the target database. In this case it is likely that all or most of the spectra will match to the same database peptide, and there is a 50% chance that this peptide will be a target. In that scenario, having observed no decoy matches, we would accept these 100 target PSMs at 1% FDR threshold, whereas the reality is that all of them are incorrect.

To demonstrate that such a failure can happen in practice, we conducted an entrapment experiment using the ISB18 data. As mentioned above, if an FDR control procedure is valid, then we expect that the FDP among its reported discoveries should not be consistently larger than the threshold. In our setting, we jointly searched the spectra from nine ISB18 runs, as an example of a dataset with moderately high spectrum multiplicity. We searched the spectra against a database containing the ISB18 proteins plus the castor plant proteome, where the castor plant proteome was considered the entrapment database, and we repeated this process 10 times with 10 different sets of decoys.

Our results suggest that TDC fails to control the PSM-level FDR in this entrapment setup. Specifically, our analysis shows that for the entire plotted range of FDR thresholds (0–10%) the FDP among the reported PSMs was consistently larger than the given threshold across all 10 searches (Figure 2A). The deviation between the FDP and the FDR threshold increased as the threshold increased. For example, at FDR thresholds of 5% and 10% the average FDP was 0.062 and 0.1148, respectively.

**Figure 2:**
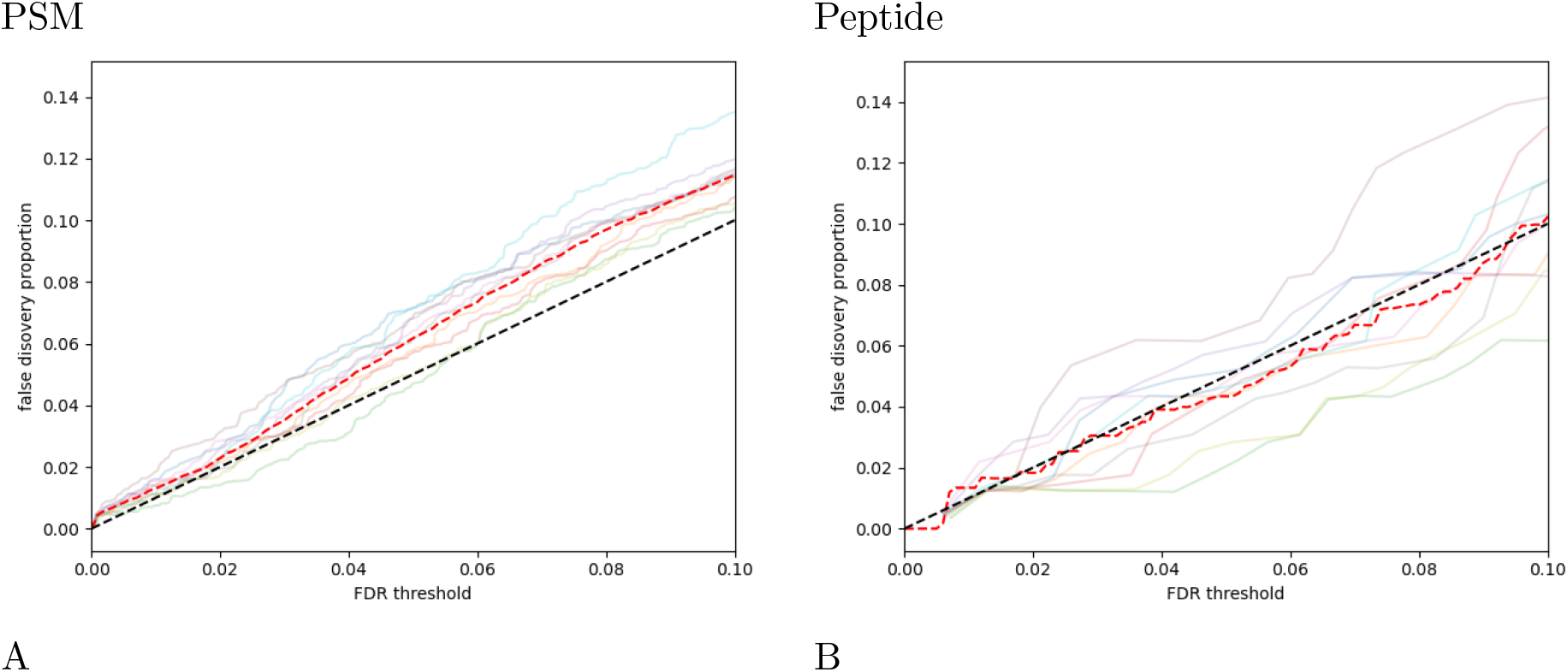
Estimated FDR when using PSM-level TDC and peptide-level TDC. The figure plots the inferred false discovery proportion (y-axis) as a function of the FDR threshold (x-axis) for a database search carried out using an entrapment setup for PSM-level (A) and peptide-level (PSM-and-peptide) (B) analysis. Each of the 10 colors corresponds to a search against a different randomly drawn decoy database, and the colors are consistent between the two plots. Notably, the average of the 10 searches (red dashed line) falls largely below the nominal FDR threshold (black dashed line) in (B) but is above it in (A). The higher variance in the peptide-level FDR plot is related to the smaller number of discoveries at each threshold: we observe an average of 3084.5 PSM-level and 156.8 peptide-level discoveries at 1% FDR.

### 3.2 Peptide-level TDC controls the FDR

To establish that the peptide-level procedures we presented here control the FDR we follow He et al.’s argument. Indeed, their peptide level analysis coincides with PSM-only, and they argued that it controls the FDR by relying on Theorem 2 of their preprint.^2^ That theorem states that assuming there is an equal chance for an incorrect identification (PSM) to be a target or a decoy match (“Equal Chance Assumption”), and assuming that this happens independently of all other identifications (“Independence Assumption”), then applying TDC (defined via (1) and (2) here) controls the FDR. They then go on to argue that it is reasonable to assume that the above two assumptions hold for the weeded-out list of PSMs, where the latter is created by removing any PSM that is matched to a peptide that has a higher-scoring PSM associated with it.

As mentioned in the Introduction applying TDC to this weeded-out list of PSMs exactly coincides with the procedure that we call “PSM-only.” Interestingly, He et al.’s argument shows that PSM-only controls the FDR at the level of filtered PSMs. In particular, this argument implies that PSM-only controls the FDR at the peptide level but it also explains why it is relatively conservative. For example, consider an incorrect PSM that involves a target peptide that is present in the sample. At the PSM level this is an incorrect identification, i.e, it is a false discovery that needs to be accounted for. However, when we switch to the peptide level, even though the PSM is incorrect, the peptide is in the sample and as such it is not a false discovery. In other words, PSM-only over-estimates the number of false discoveries, which is consistent with our observations below that this procedure typically reports significantly fewer peptides at any given threshold than the peptide-only and PSM-and-peptide methods.

The above approach naturally extends to arguing that peptide-only, as well as PSM-and-peptide, controls the FDR. Indeed, first note that Theorem 2 applies more generally to a list of hypotheses such that with each hypothesis we associate a target/decoy (win) label as well as a score. In this context the Equal Chance Assumption means that for each true null hypothesis the label is equally likely to be a target or decoy, and the Independence Assumption means that this should happen independently of all the scores and all the other labels. This extension follows immediately from the proof of He et al., or alternatively this is Theorem 3 of^3^ applied to Selective SeqStep+ with *c* = 1/2.

In applying this extension to peptide-only and PSM-and-peptide, our hypotheses essentially coincide with the list of target peptides, more specifically, each null hypothesis is that the corresponding peptide is *not* in the sample. For a true null hypothesis, that is, for an out-of-sample peptide, it is reasonable to assume that its corresponding decoy is equally likely to win the head-to-head competition between the two. Moreover, we argue, similarly to He et al., that it is reasonable to assume that this happens independently of all the scores, as well as all other labels (a point we will briefly revisit in the Discussion).

Finally, to demonstrate empirically that these procedures apparently control the FDR we conducted the same entrapment analysis we did for the PSM-level TDC. Figure 2B suggests that PSM-and-peptide controls the peptide-level FDR because the peptide-level FDP values center around the *y* = *x* line across the entire plotted threshold range. Note that our procedures are designed to control the FDR, which is the expected value of the FDP. Hence, it is not surprising that, as observed, the FDP can exceed the prescribed thresholds. Notably, the average of the FDP across the 10 searches (dashed red line) is largely below nominal FDR threshold (dashed black line) for the peptide-level TDC of PSM-and-peptide while it is above the line for the PSM-level analysis. The results for PSM-only and peptide-only are essentially the same (and further empirical validation of PSM-only can be found in^2^).

### 3.3 Peptide-level FDR control benefits from peptide-level competition

Next, we focused on comparing the statistical power of our three different methods for estimating peptide-level FDR: PSM-only, peptide-only, and PSM-and-peptide. For this analysis, we searched runs from castor plant, *E. coli*, human, mouse, and yeast against their respective databases, and we counted the number of reported discoveries at various FDR thresholds (0–10%) for each of the three FDR procedures. Each run was searched four times against the same protein database but with a different score function each time (XCorr, XCorr p-value, Tailor, and combined p-value).

Empirically, we found that PSM-and-peptide yielded the best performance across all runs and score functions (Figure 3). Conversely, we found that the PSM-only procedure had the overall worst performance, while the peptide-only procedure always had middling performance. For example, in the *E. coli* run at a 1% FDR, PSM-and-peptide outperformed PSM-only by 1050 (11.33%), 2397 (25.85%), 3040 (41.02%), and 881 (10.38%) peptide detections for the combined p-value, Tailor, XCorr, and XCorr p-value score functions, respectively. Similarly, at the same 1% threshold PSM-and-peptide outperformed peptide-only by 589 (5.84%), 544 (5.35%), 1338 (14.68%), and 364 (4.04%) peptide detections. An analysis of a second run from each of the five datasets showed similar results (Supplementary Figure 1).

**Figure 3:**
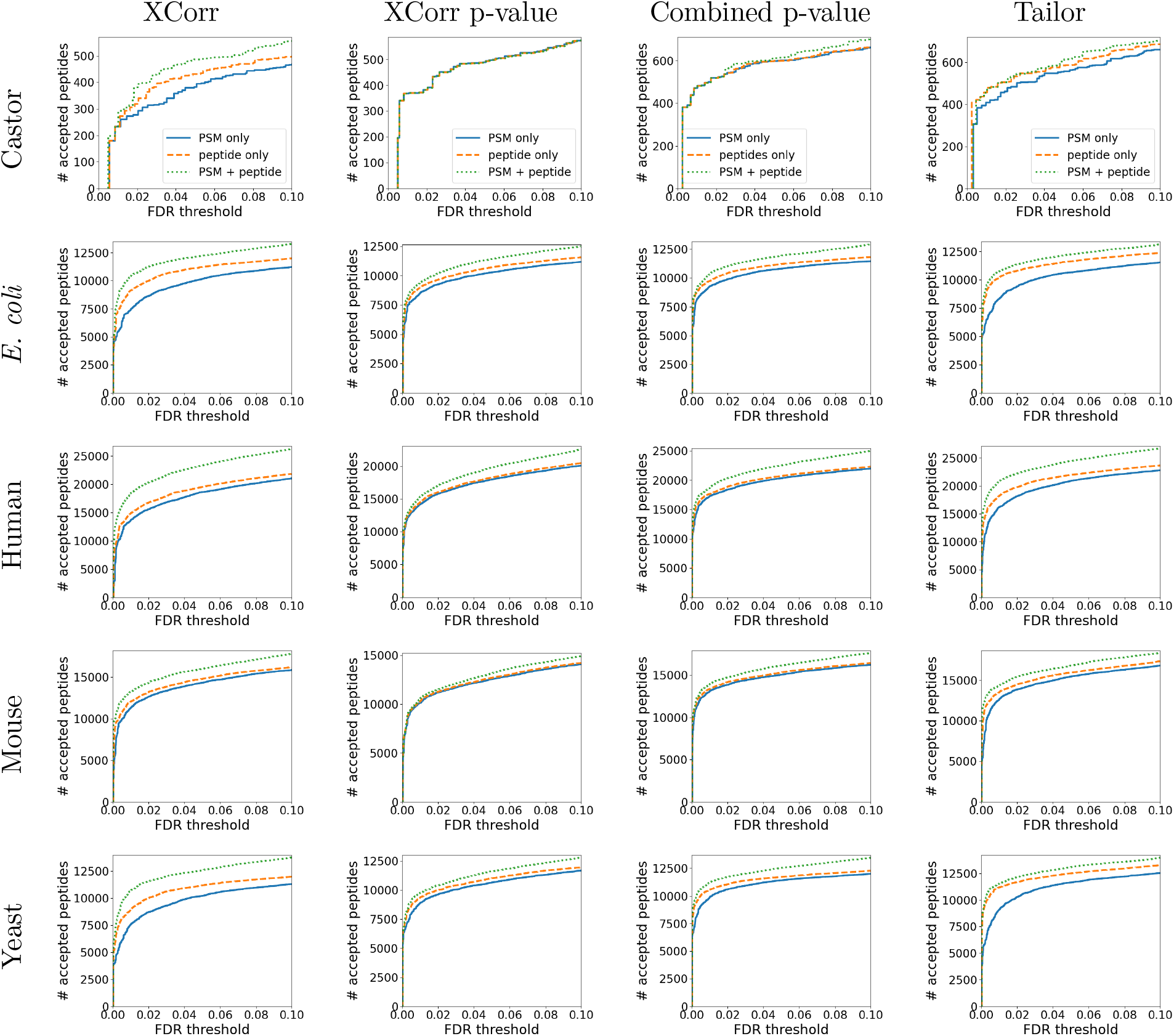
Peptide-level FDR procedures. Each plot compares the performance of three different peptide-level FDR estimation and control procedures. Each row of panels represents a run from a different species, and each column represents a different score function (XCorr, XCorr p-value, combined p-value, and Tailor score). The PSM-and-peptide method generally has the best performance for all score functions and runs across the plotted range of FDR thresholds (0–10%). Supplementary Figure 1 shows similar results using a different run from each of the five datasets.

We observed that the performance boost of PSM-and-peptide, when compared to peptide-only and PSM-only, was generally greatest when using a non-calibrated score. In this setting, a score is calibrated if a particular value *X* has the same significance regardless of the peptide that was scored.^26^ Specifically, we noted that in nine out of 10 runs, the performance boost when using PSM-and-peptide was greatest when using the non-calibrated XCorr score. The one exception was a castor plant run where the heuristically-calibrated Tailor score had the greatest boost in performance instead of the XCorr score. This observation is not surprising, because the additional peptide-level competition in PSM-and-peptide confers a form of calibration.

## 4 Discussion

We argued here, as did He et al. before us,^2^ that multiplicity, where multiple spectra are generated by the same peptide species, can impair TDC’s ability to control the PSM-level FDR. To avoid this problem, we therefore suggest that mass spectrometrists should employ FDR control exclusively at the peptide or protein levels.

An alternative way to circumvent the problem with PSM-level FDR control is to cluster the spectra in the hope that each cluster of spectra corresponds to a single peptide species.^27^ However, it is not clear whether this approach—with its added complexity of how to define the clusters—offers any advantage over simply switching to peptide-level analysis in the first place.

One question that intrigued us is whether we can use the multiplicity to improve our peptide-level analysis. Specifically, we postulated that if, in addition to its score, we assign to each peptide its multiplicity, i.e., the number of PSMs that were optimally matched to the peptide, then we could better distinguish between peptides that were truly present in the samples and false detections. To test this hypothesis, we used our recently developed Group-walk^28^ to divide the peptides into groups according to their multiplicity and compared that approach to a single-group approach that ignores the multiplicity. While we did see a fairly consistent power gain when using the multiplicity in this way, that gain was unfortunately marginal (about 0.5% at 1% FDR), so we concluded that it does not justify the added complexity of the procedure.

We already discussed above why the commonly used method for estimating peptide-level FDR using the PSM-only approach is conservative. To see why the peptide-only approach is more conservative note that both the target and decoy peptide scores are higher for this procedure than they are for PSM-and-peptide: each spectrum generates two PSM scores in peptide-only but only one PSM score in PSM-and-peptide and, in both cases, each peptide score is defined as the maximum of all PSMs that include that peptide. Now consider a target peptide that is in the sample. Assuming that the peptide generates one or more spectra, then regardless of whether we use peptide-only or PSM-and-peptide, this peptide’s score is most likely to be defined by one of the corresponding PSMs involving those generated spectra. Therefore, in general, the score of a peptide that is present in the sample does not increase in the concatenated search of PSM-and-peptide. At the same time, the corresponding decoy score will generally increase as part of the general trend of increasing scores, thus offering a tougher competition to the in-sample target peptide.

One practical challenge in switching from the PSM-only to the PSM-and-peptide procedure is the requirement that the search engine retains the pairings between each target and its shuffled decoy peptide. This pairing information must be retained during the creation of the decoy database and reported to the user for use during FDR control. The Tide search engine supports this type of analysis by reporting, for each decoy PSM, the corresponding target peptide and by optionally generating a list of targets and corresponding decoys during the database indexing step.

In this work, we focused on TDC for two reasons. First, TDC is the most commonly used approach to controlling the FDR in the analysis of mass spectrometry data. Second, it is a fairly flexible method that, in principle, can work with any score function as long as competing decoy scores can be computed. However, there are alternative approaches that have been suggested in the literature, including relying on canonical p-value based FDR controlling procedures such as Benjamini and Hochberg’s procedure.^29^ The p-values in this case can be taken directly from the search tool, if it provides those, or alternatively estimated from decoy scores, either using a dedicated decoy database or a generic/universal one.^30–32^ Bayesian FDR analysis offers a different route, where one models the scores using a two-component mixture distribution as pioneered by PeptideProphet,^33^ with a more recent model using a skew-normal rather than a Gamma distribution.^34^

Finally, He et al. posited that their Equal Chance Assumption—that an incorrect identification is equally likely to be a target or a decoy peptide—applies to their peptide-level analysis. The same rationale applies to the other two methods presented here, but there is one caveat to this assumption (that applies in all these cases), namely, that the assumption is questionable when the target database contains a non-trivial proportion of close neighbors (peptides whose theoretical spectra are highly similar to one another^35^). Fortunately, in practice this does not seem to be a common problem, and this is something that could be addressed in the future. Keep in mind, though, that the existence of the “neighbors” phenomenon complicates any approach to FDR control, and its impact is not limited to TDC (although the latter has its idiosyncratic limitations when, unlike our analysis here, it is applied to scores produced by a post-processor^36^).

## Acknowledgments

Some of research described in this paper was conducted under the Laboratory Directed Research and Development Program at Pacific Northwest National Laboratory, a multiprogram national laboratory operated by Battelle for the U.S. Department of Energy. Andy Lin is grateful for the support of the Linus Pauling Distinguished Postdoctoral Fellowship program. Pacific Northwest National Laboratory is a multiprogram national laboratory operated by Battelle Memorial Institute for the United States Department of Energy under contract DE-AC06-76RLO. This work was funded in part by National Institutes of Health award R01 GM121818.

We are also grateful to the anonymous referees for their comments and suggestions which helped improve this manuscript.

## 5 Supporting Information

- **Supplemental File S1:** PDF containing Supplementary Information.

## For Table of Contents Only

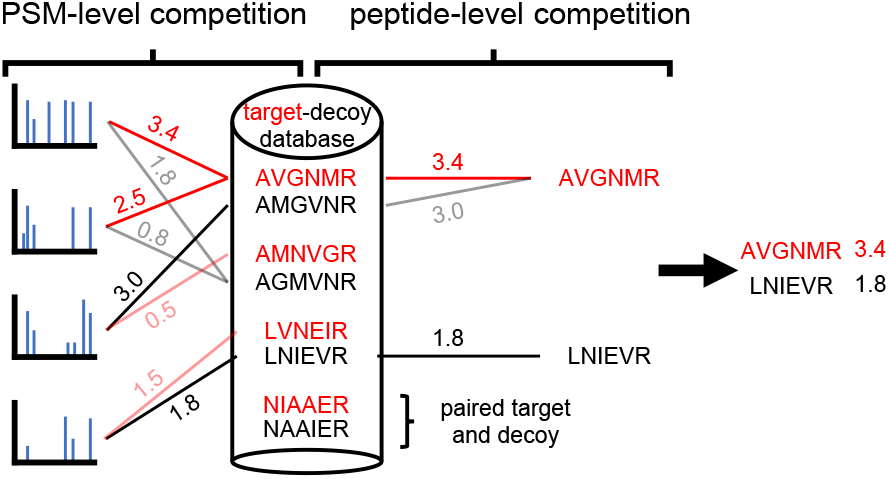

