## Supplemental File for "Improving peptide-level mass spectrometry analysis via double competition"

### Supplement to "Detecting more peptides from bottom-up mass spectrometry data via peptide-level target-decoy competition"

Andy Lin[1], Temana Short[2], William Stafford Noble[3,4], and Uri Keich[2]

[1]Chemical and Biological Signatures, Pacific Northwest National Laboratory, Seattle, WA, USA 98109
[2]School of Mathematics & Statistics, University of Sydney, NSW 2006, Australia
[3]Department of Genome Sciences, University of Washington, Seattle, WA, USA 98195
[4]Paul G. Allen School of Computer Science and Engineering, University of Washington, Seattle, WA, USA 98195

September 26, 2022

#### Supporting Information

- **Supplemental File S1:** PDF containing Supplementary Information.

| Variable | Definition |
|---|---|
| $S$ | spectra |
| $\mathcal{T}$ | target database |
| $\mathcal{D}$ | decoy database |
| $TD\_labels$ | denotes whether a peptide is a target or decoy: (peptide_id, is_decoy) |
| $TD\_pairing$ | denotes paired target and decoy: (target_id, decoy_id) |
| $PSMs$ | data structure of triplets per PSM: (spectrum_id, peptide_id, score) |
| $PS$ | peptide scores; (peptide_id, score) |
| $\alpha$ | user defined FDR threshold |
| $A$ | set of matches that pass FDR threshold ($a_i = 1$ is the ith peptide's score above the TDC threshold) |
| $R$ | set of reported matches |

Table 1: **Notation.** Notation used in this supplement.

**Algorithm 1 PSM-only**

**procedure** PSM-ONLY($S, \mathcal{T}, \mathcal{D}, \alpha$) ▷ $\mathcal{D}$ generated from $\mathcal{T}$
  $(PSMs)$ := SEARCH($S, \mathcal{T} \cup \mathcal{D}$) ▷ yields top database match per spectrum
  $(PS)$ := BESTSCOREPERPEPTIDESEQUENCE($PSMs$)
  $(TD\_labels)$ := ASSIGNTDLABELS($PS, T, D$)
  $A$ := CONTROLFDR($PS$, $TD\_labels$, $\alpha$) ▷ implements Equations (1) and (2)
  $R := \{(p_i, PS(p_i)) \mid a_i = 1, p_i \in \mathcal{T}\}$
  **return** R
**end procedure**

**Algorithm 2 peptide-only**

**procedure** PEPTIDE-ONLY($S, \mathcal{T}, \mathcal{D}, TD\_pairing, \alpha$) ▷ $\mathcal{D}$ generated from $\mathcal{T}$
  $target\_PSMs$ := SEARCH($S, \mathcal{T}$)
  $decoy\_PSMs$ := SEARCH($S, \mathcal{D}$)
  $target\_PS$ := BESTSCOREPERPEPTIDESEQUENCE($target\_PSMs$)
  $decoy\_PS$ := BESTSCOREPERPEPTIDESEQUENCE($decoy\_PSMs$)
  $(PS, TD\_labels)$ := COMPETEPAIREDTARGETANDDECOY($target\_PS \cup decoy\_PS, TD\_pairing$)
  $A$ := CONTROLFDR($PS$, $TD\_labels$, $\alpha$) ▷ implements Equations (1) and (2)
  $R := \{(p_i, PS(p_i)) \mid a_i = 1, p_i \in \mathcal{T}\}$
  **return** R
**end procedure**

**Algorithm 3 PSM-and-peptide**

**procedure** PSM-AND-PEPTIDE($S, \mathcal{T}, \mathcal{D}, TD\_pairing, \alpha$) ▷ $\mathcal{D}$ generated from $\mathcal{T}$
  $(PSMs)$ := SEARCH($S, \mathcal{T} \cup \mathcal{D}$) ▷ yields top database match per spectrum
  $(PS)$ := BESTSCOREPERPEPTIDESEQUENCE($PSMs$)
  $(PS_1, TD\_labels_1)$ := COMPETEPAIREDTARGETANDDECOY($PS, TD\_pairing$)
  $A$ := CONTROLFDR($PS_1$, $TD\_labels_1$, $\alpha$) ▷ implements Equations (1) and (2)
  $R := \{(p_i, PS(p_i)) \mid a_i = 1, p_i \in \mathcal{T}\}$
  **return** R
**end procedure**

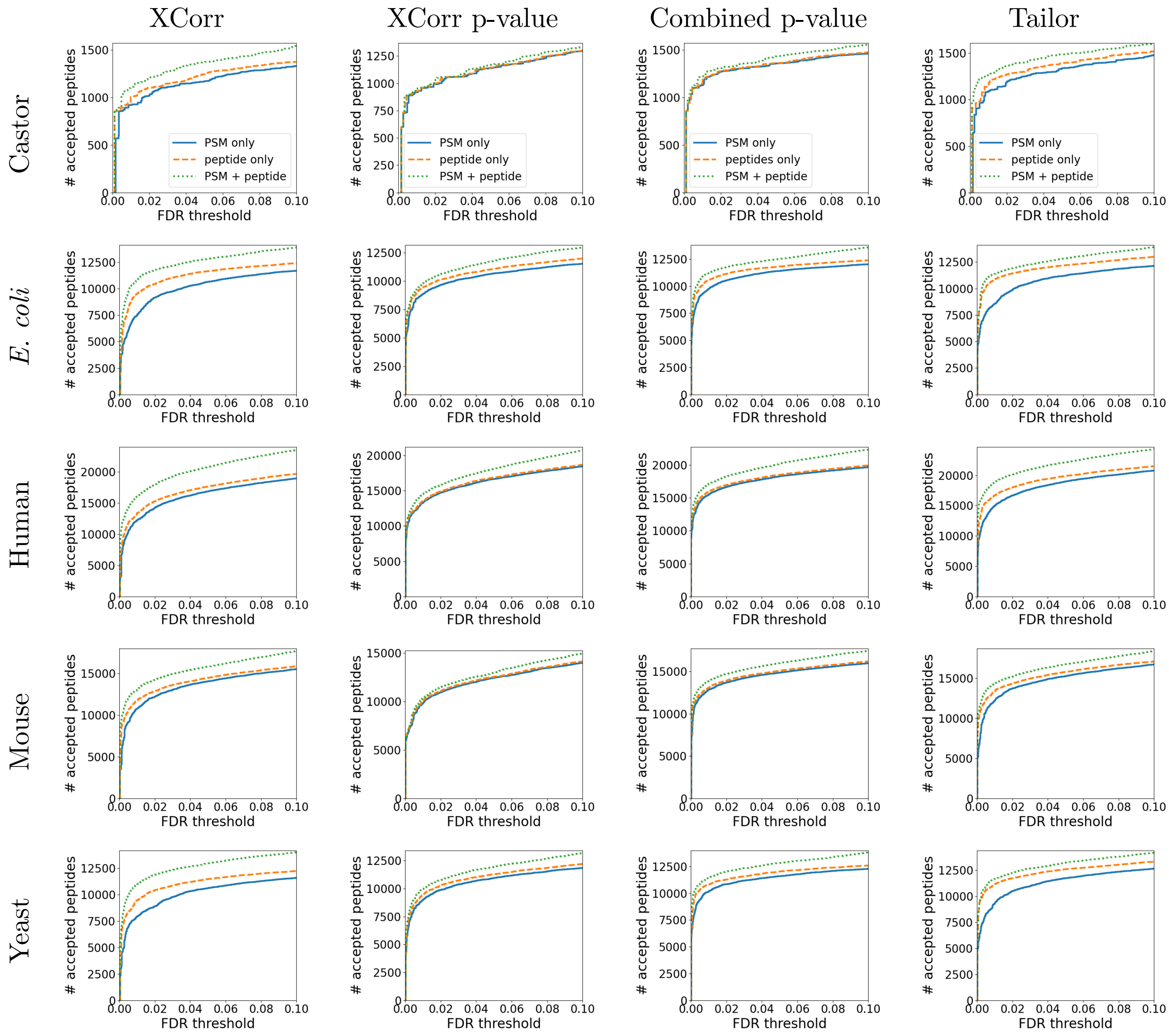


Figure 1: **Peptide-level FDR procedures.** Each plot compares the performance of three different peptide-level FDR estimation procedures. Each row of panels represents a different score function (XCorr, XCorr p-value, combined p-value, and Tailor score). Each column is a run from a different species. The PFPC method generally has the best performance for all score functions and runs across the plotted q-value range of 0–10%.
